# A gatekeeping role of ESR2 to maintain the primordial follicle reserve

**DOI:** 10.1101/2020.02.06.937953

**Authors:** V. Praveen Chakravarthi, Subhra Ghosh, Katherine F. Roby, Michael W. Wolfe, M. A. Karim Rumi

**Author notes:** **Correspondence:** M. A. Karim Rumi.

## Abstract

Over the entire reproductive lifespan in mammals, a fixed number of primordial follicles serve as the source of mature oocytes. Uncontrolled and excessive activation of primordial follicles can lead to depletion of the ovarian reserve. We observed that disruption of ESR2-signaling results in increased activation of primordial follicles in *Esr2*-null (*Esr2-/-*) rats. However, follicle assembly was unaffected, and the total number of follicles remained comparable between neonatal wildtype and *Esr2-/-* ovaries. While the activated follicle counts were increased in *Esr2-/-* ovary, the number of primordial follicles were markedly decreased. Excessive recruitment of primordial follicles led to premature ovarian senescence in *Esr2-/-* rats and was associated with reduced levels of serum AMH and estradiol. Disruption of ESR2-signaling through administration of a selective antagonist (PHTPP) increased the number of activated follicles in wildtype rats, whereas a selective agonist (DPN) decreased follicle activation. In contrast, primordial follicle activation was not increased in the absence of ESR1 indicating that the regulation of primordial follicle activation is ESR2-specific. Follicle activation was also increased in *Esr2*-mutants lacking the DNA-binding domain, suggesting a role for the canonical transcriptional activation function. Both primordial and activated follicles express ESR2 suggesting a direct regulatory role for ESR2 within these follicles. We also detected that loss of ESR2 augmented the activation of AKT, ERK and mTOR pathways. Our results indicate that the lack of ESR2 upregulated both granulosa and oocyte factors, which can facilitate AKT and mTOR activation in *Esr2-/-* ovaries leading to increased activation of primordial follicles.

## Introduction

Primordial follicles generated during early life serve as the source of fertilizable oocytes in mammals throughout the reproductive lifespan (1). A limited number of primordial follicles are steadily recruited into the growing pool while the remaining are maintained in the dormant state (2,3). The female reproductive life comes to an end when folliculogenesis ceases due to the loss of primordial follicles (4,5). Regulation of the dormancy and selective activation of primordial follicles plays an important role in determining the reproductive lifespan in females (6). An uncontrolled and increased activation of primordial follicles leads to excessive depletion of the ovarian reserve and development of premature ovarian insufficiency (POI), which causes infertility and estrogen deficiency in 1-2% of all women of reproductive age (7–9). Pathological conditions as well as certain therapeutic procedures that deplete the follicle reserve can also lead to POI (10). Understanding the regulatory mechanisms of primordial follicle activation remains critical for prevention or treatment of POI.

Activation of primordial follicles to primary follicles is a strictly regulated and steady process. This activation is gonadotropin independent, but the exact molecular mechanisms of the prolonged dormancy and selective recruitment of primordial follicles remain largely unclear (11). Primordial follicles consist of dormant oocytes surrounded by a few flattened granulosa cells (GCs). Activation of the primordial follicle is characterized by growth and differentiation of the flattened GCs to cuboidal GCs as well as oocyte growth and deposition of the zona pellucida (ZP) matrix by the oocytes (12–14). Signaling pathways in both GCs and oocytes have been found to play important roles in primordial follicle activation (15). It has been suggested that the process starts in the GCs, and the signals from GCs subsequently activating the oocytes in primordial follicles (16,17). Conversely, studies have also indicated that the activation process can be initiated in the oocytes of primordial follicles (14). Inhibitory factors secreted from the primordial follicles or from activated follicles may also maintain the primordial follicles in a dormant state (18).

Previous studies with mutant mouse models have identified several growth factors and cytokines involved in primordial follicle activation. While TGF-β superfamily members such as bone morphogenic proteins (BMPs) and growth and differentiation factors (GDFs) act as positive regulators (19,20), AMH acts as a negative regulator of primordial follicle activation (21). Growth factors unrelated to the TGF-β superfamily including KITLG (22) and FGF2 (23), also enhance the activation process. These growth factors and cytokines regulate key signaling pathways and transcriptional regulators. Regulation of PI3 kinase (PI3K) and mTOR-signaling and the downstream transcription factor FOXO3A are essential regulators of primordial follicle activation (14,24). In addition to FOXO3A (25), several other transcription factors have been found to be important for the regulation of follicle activation. FOXL2 (26) in primordial GCs as well as SOHLH1 (27), SOHLH2 (28), NOBOX (29), LHX8 (27), GATA4, and GATA6 (30) in oocytes carry out important regulatory functions. We recently observed that transcriptional regulation mediated by estrogen receptor β (ESR2) plays an essential role during primordial follicle activation.

ESR2 is the predominant estrogen receptor in the ovary (31–34). ESR2 polymorphisms and mutations in women have been linked to ovulatory dysfunctions, including complete ovarian failure (35–38). *Esr2*-mutant mouse (39,40) and rat models (41) suffer from defective follicle development and failure of ovulation. Studies have suggested that estrogen signaling plays a role in follicle assembly and early follicle development (42,43), but a role of ESR2 in ovarian primordial follicle activation has not yet been demonstrated. In this study, we observed that disruption of ESR2 signaling markedly increases the activation of primordial follicles. Our findings suggest that the transcriptional regulatory function of ESR2 plays a gatekeeping role in maintaining the primordial follicle reserve.

## Materials and Methods

### Animal models

Wildtype, *Esr1-*knockout, and *Esr2-mutant* Holtzman Sprague-Dawley (HSD) rats were included in this study. The *Esr1*-knockout rat model was generated by the deletion of exon 3 in the *Esr1* gene, which caused a frameshift and null mutation (*Esr1-/-*) (44). *Esr2*-mutant rat models were generated by targeted deletion of the exon 3 or exon 4 in the *Esr2* gene (41). Exon 3 deletion caused a frameshift and null mutation (*Esr2-/-*), whereas exon 4 deletion resulted in a mutant ESR2 protein lacking the DNA binding domain (DBD). Rats were screened for the presence of mutations by PCR using tail-tip DNA samples (RED extract-N-Amp Tissue PCR Kit, Sigma-Aldrich) as previously described (41,44). All procedures were performed in accordance with the protocols approved by the University of Kansas Medical Center Animal Care and Use Committee.

### Gonadotropin treatment for follicle maturation

28-day-old *Esr2*-/- and age-matched wildtype female rats were used for the evaluation of gonadotropin-induced ovarian follicle development. Synchronized follicle maturation was induced by intraperitoneal injection of 30 IU PMSG (Lee Biosolutions, Maryland Heights, MO). 48h after the PMSG injection, 30 IU of hCG (Lee Biosolutions) was injected intraperitoneally. 4h after hCG injection, ovaries were collected and processed for histological examination, or cumulus-oocyte complexes (COCs) were harvested by needle-puncture of the large antral follicles under microscopic examination as described previously (45). Cumulus cells were mechanically detached from the COCs by repeated pipetting, and the oocytes were counted under microscopic examination.

### Treatment with selective ESR2-antagonist and -agonist

A selective ESR2 antagonist PHTPP (46) or a selective ESR2 agonist DPN (47) (Cayman Chemical, Ann Arbor, MI) was dissolved in DMSO (5µg/µl). 2µl of PHTPP (10µg) or 2µl of DPN (10µg) was further diluted in 50µl of sesame oil (Sigma-Aldrich, St. Louis, MO) and injected subcutaneously into wildtype rats from postnatal day (PND) 5 to 15. Rats were sacrificed on PND16, and the ovaries were processed either for histological examination or for total RNA and protein analyses.

### Blood collection and hormone assays

Blood samples were collected by cardiac puncture at sacrifice. After clotting at room temperature, serum was separated by centrifugation and frozen at −80C. Serum AMH level was determined by an enzyme immunoassay (AL-113, Ansh Labs, Webster, TX). Total serum 17β-estradiol (E2) and serum progesterone (P4) levels were measured by RIA as described previously (48).

### Total follicle counting in the serial sections of whole ovary

Ovaries were collected and fixed in 4% formaldehyde overnight, processed, and embedded in paraffin following standard procedures (49–51). Whole ovaries were serially sectioned at 6 µM thickness and stained with hematoxylin and eosin (H&E) (49–51). The stages of follicle development including primordial, primary, secondary, early antral, and antral were determined as described previously (52). The primordial follicles were recognized as small oocytes surrounded by a few flattened GCs, the primary follicles contained larger oocytes surrounded by a single layer of cuboidal GCs, and the oocytes in the secondary follicles were surrounded by multiple layers of GCs. The tertiary follicles were categorized into early antral and antral follicles based on appearance and extent of the cavities within the follicles (Figure 1D). Atretic follicles were recognized by pyknotic GCs surrounding degenerated oocytes. The follicles were counted on every fifth section under light microscopy (49,50). To avoid counting the same follicle more than once, primordial and primary follicles were counted if they exhibited a nucleus, whereas the secondary, early antral, and antral follicles were counted only in presence of a nuclei with prominent nucleoli (49–51). The counts were multiplied by 5 to obtain the total count of follicles in the whole ovary.

**Figure 1.**
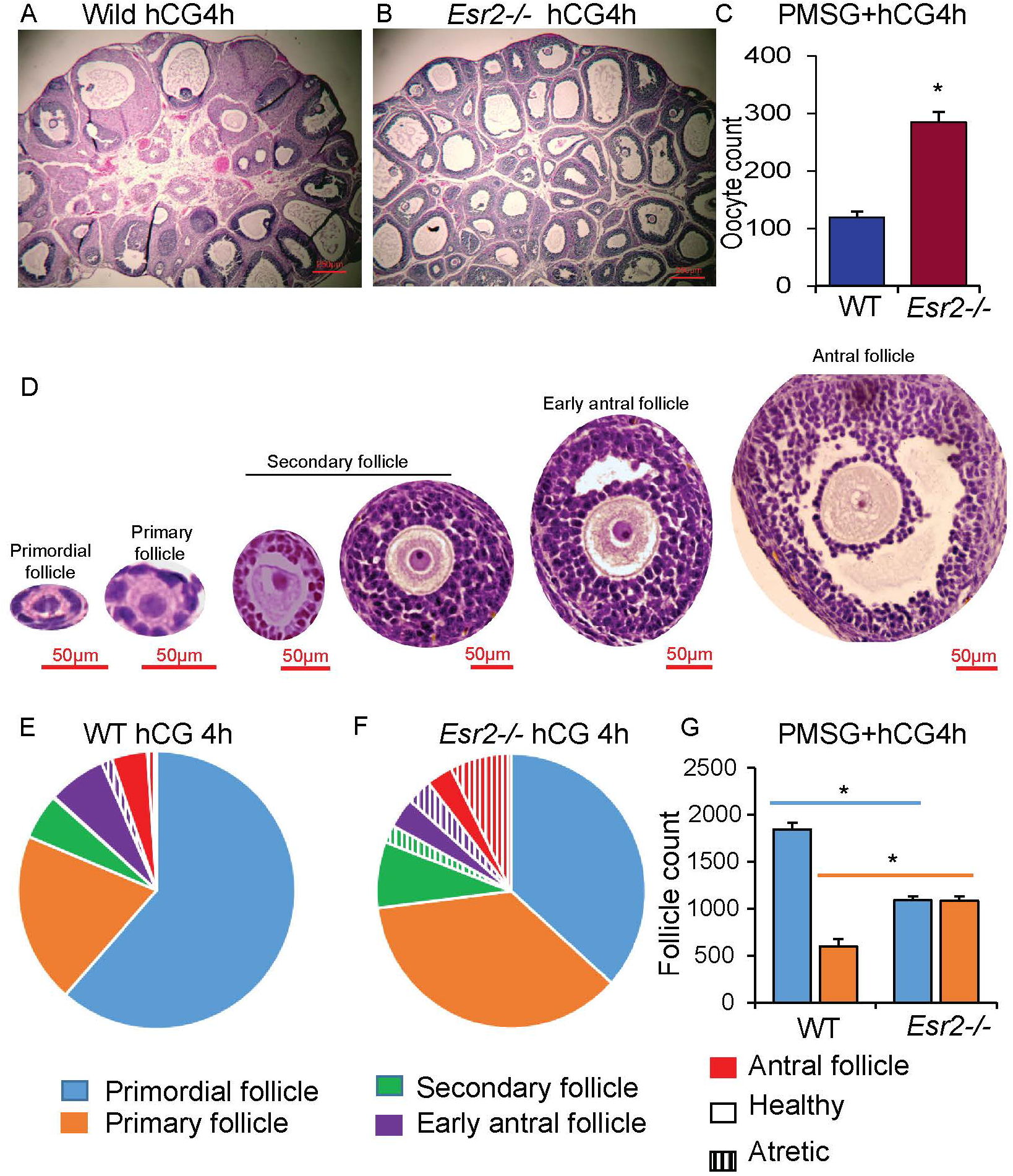
Gonadotropin stimulation resulted in an increased number of antral follicles in *Esr2-/-* ovaries. Wildtype (WT) and *Esr2-/-* rats were treated with exogenous gonadotropins on PND28. 48h after PMSG (30IU) injection, rats were administered with hCG (30IU). 4h after the hCG injection, ovaries were collected and processed for histological examination or collection of COCs. Histological examination demonstrated the presence of ovarian follicles at different stages of development in WT rats (**A**). In contrast, *Esr2-/-* rat ovaries showed an increased number of antral follicles (**B**). Cumulus cells were detached from the oocytes by mechanical pipetting before counting under microscope. Oocyte yield was about three-fold higher in *Esr2-/-* rat ovaries (**C**). Serially sectioned whole ovaries were stained with H&E, and follicles at different stages were counted in WT and *Esr2-/-* rats (**D**). Follicle counting demonstrated a decreased number of primordial follicles and an increased number of activated follicles in *Esr2-/-* rats (**E-G**). Data shown as mean ± SE, n≥3. \**P* ≤ 0.05.

### Protein extraction and western blotting

Ovaries were collected on PND 4, 6, and 8, and total protein was extracted from the ovaries using 1X SDS lysis buffer (62.5mMTris-HCl pH 6.8, 2% SDS, 42 mM dithiothreitol, 10% glycerol, and 0.01% bromophenol blue), containing protease and phosphatase inhibitors (Cell Signaling Technologies, Danvers, MA). Ovarian lysates were sonicated to shear DNA and reduce viscosity, heat denatured, and separated on a 4-20% SDS-PAGE. Electrophoresed proteins were transferred from the gel to PVDF membranes, blocked with 5% skim milk in TBST (1xTBS buffer containing 0.1% Tween-20), and incubated for 1h at room temperature with specific primary antibodies (Table 1) at the appropriate dilution in blocking solution. After removing the unbound primary antibody solution, membranes were washed with TBST, blocked, and incubated with peroxidase-conjugated anti-mouse, or anti-rabbit secondary antibodies (Jackson Immunoresearch, West Grove, PA) at a dilution of 1:25,000 to 50,000, and the immunoreactivity signals were visualized with Luminata Crescendo HRP substrate (Millipore Sigma, Burlington, MA).

**Table 1.** List of antibodies used in the western blot assays

| Target | Name of Antibody | Manufacturer, Catalog number | Species, Mono or Polyclonal | Dilution | RRID |
| --- | --- | --- | --- | --- | --- |
| ESR2 | Anti-ESR2, clone 68-4 | EMD Millipore, 05-824 P | Rabbit; monoclonal | 5000 | AB_11212759 |
| pAKT | Anti-Phospho-Akt (T308) | Cell Signaling Technology, 13038 | Rabbit; monoclonal | 2000 | AB_2629447 |
| pAKT | Anti-Phospho-Akt (Ser473) | Cell Signaling Technology, 4060 | Rabbit; monoclonal | 2000 | AB_2315049 |
| AKT | Anti- Akt | Cell Signaling Technology, 4691 | Rabbit; monoclonal | 2000 | AB_915783 |
| PTEN | PTEN (D4.3) | Cell Signaling Technology, 9188 | Rabbit; monoclonal | 2000 | AB_2253290 |
| ERK | Anti-p44/42 MAPK (ERK1/2) | Cell Signaling Technology, 4695 | Rabbit; monoclonal | 1000 | AB_390779 |
| pERK | Anti-Phospho-ERK1-T202/Y204 + ERK2-T185/Y187 | Cell Signaling Technology, 4370 | Rabbit; monoclonal | 2000 | AB_2315112 |
| pTSC2 | Anti-Phospho-TSC2-T1462 | ABclonal Technology, AP0866 | Rabbit; polyclonal | 2000 | AB_2771631 |
| TSC2 | Anti-TSC2 | ABclonal Technology, A0492 | Rabbit; polyclonal | 2000 | AB_2757221 |
| pmTORC1 | Anti-Phospho-mTOR (Ser2448) | Cell Signaling Technology, 5536 | Rabbit; monoclonal | 2000 | AB_10691552 |
| mTORC1 | Anti-mTOR | Cell Signaling Technology, 2983 | Rabbit; monoclonal | 2000 | AB_2105622 |
| pP70S6K | Anti-pP70S6Kinase (T389) | Cell Signaling Technology, 9234 | Rabbit; monoclonal | 2000 | AB_2269803 |
| pRPS6 | Anti-pS6 Ribosomal Protein (Ser240/244) | Cell Signaling Technology, 5364 | Rabbit; monoclonal | 2000 | AB_10694233 |
| ACTB | Anti-β-actin | Sigma-Aldrich (AC15), A 1978 | Mouse; monoclonal | 25,000 | AB_476692 |

### RNA extraction and RT-qPCR

Total RNA was extracted from the PND8 ovaries using TRI Reagent (Sigma-Aldrich). 1000ng of total RNA from each sample was used for the preparation of cDNA using High-Capacity cDNA Reverse Transcription (RT) Kits (Applied Biosystems, Foster City, CA). The cDNA was diluted 1:50 in 10mM Tris-HCl (pH 7.4) and 2.5 µl of the diluted cDNA was used in a 10 µl quantitative PCR (qPCR) reaction mixture containing Applied Biosystems Power SYBR Green PCR Master Mix (Thermo Fisher Scientific). Amplification and fluorescence detection of RT-qPCR were carried out on Applied Biosystems QuantStudio Flex 7 Real Time PCR System (Thermo Fisher Scientific). The ΔΔCT method was used for relative quantification of target mRNA expression and normalized to Rn18s (18S rRNA) level. A list of qPCR primers is shown in Table 2.

**Table 2.**
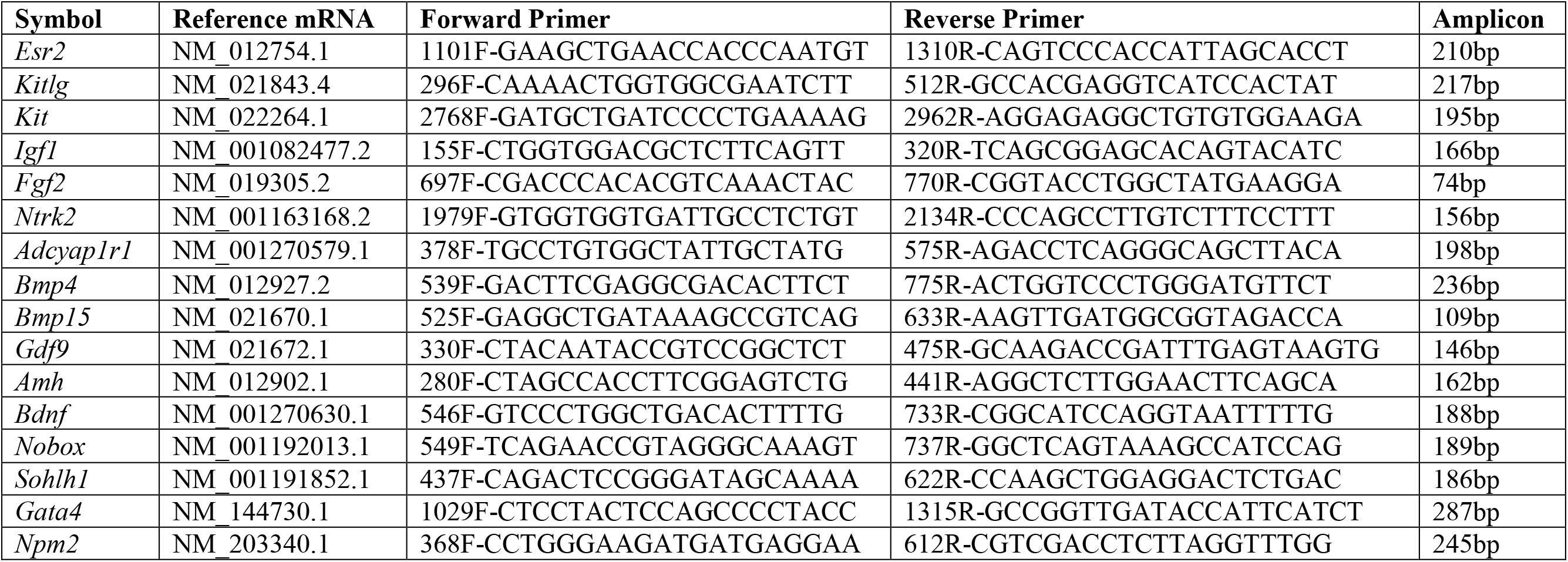
List of primers used in the qPCR assays

### Isolation of primordial and primary follicles

Approximately 100 mg of minced PND12 rat ovary tissue was digested in 1ml of digestion medium [199 media containing 0.08 mg/ml of liberase with medium-concentration of thermolysin (Roche Diagnostics GmbH, Mannheim, Germany) supplemented with 5U/ml of DNase I and 1% BSA (Thermo Fisher Scientific, Waltham, MA)]. The digestion mix was agitated on an orbital shaker (Disruptor Genie, Scientific Industries, Bohemia, NY) at 1500 rpm for 30 min at room temperature. The enzymatic reaction was stopped by addition of 10% FBS. Digested ovary tissues were passed through a 70µM cell strainer (Thermo Fisher Scientific) to remove the secondary, and large follicles as well as tissue aggregates. The filtrate containing the small follicles and cellular components was filtered again through a 35µM cell strainer (BD Falcon, Franklin Lakes, NJ). The 35µM strainer was reverse eluted with medium 199 to isolate the primary follicles and the filtrate was subjected to sieving through a 10µM cell strainer (PluriSelect USA, Gillespie Way, CA) to separate the primordial follicles from other cellular components. Finally, the 10µM cell strainer was reverse eluted to isolate the primordial follicles. Unwanted cellular components were removed from the desired follicles under microscopic examination before proceeding to RNA isolation.

### Gene expression analyses in primordial and primary follicles

We used 200-250 primordial follicles and 100-150 primary follicles for cDNA synthesis using the Message Booster cDNA synthesis kit (Lucigen, Palo Alto, CA). Direct cDNA and subsequent cRNA syntheses were performed by following the manufacturer’s instruction. *In vitro* synthesized cRNA was purified by using Monarch RNA cleanup kit (New England Biolabs, Ipswich, MA) and subjected to first strand, and subsequent second strand cDNA synthesis using the regents provided in the Message Booster cDNA synthesis kit. The cDNA was diluted 1:10 in 10mM Tris-HCl (pH 7.4) and 2.5 µl of the diluted cDNA was used in 10 µl qPCR reaction as described above. The relative quantification of target mRNA expression was calculated by normalizing the data with *Actb* expression.

### Statistical analyses

Follicle counting was performed on more than 3 individual rats of the same genotype at each time point. Gene expression analyses were performed on at least 6 individual rats. The experimental results are expressed as mean ± SE. Statistical comparisons between two means were determined with Student’s t-test while comparisons among multiple means were evaluated with ANOVA followed by Duncan post hoc test. *P* values ≤ 0.05 were considered as significant level of difference. All statistical calculations were done with SPSS 22 (IBM, Armonk, NY).

## Results

### Increased number of antral follicles in *Esr2-/-* rats

Exogenous gonadotropins were administered to PND28 prepubertal rats to induce follicle maturation. Compared to the wildtype, *Esr2-/-* ovaries exhibited a larger number of antral follicles 4h after hCG administration into PMSG primed rats (Figure 1A, B). At this time point, cumulus-oocyte complexes (COCs) were isolated from the gonadotropin-stimulated ovaries by needle puncture, and cumulus cells were mechanically removed to count the oocytes. Oocyte counts were significantly higher in *Esr2*-/- rats (Figure 1C). To compare the extent of follicle maturation between wildtype and *Esr2-/-* rats, total follicle counting was performed in serial sections of gonadotropin-stimulated ovaries. While the number of primordial follicles were decreased, the number of primary, secondary, and antral follicles were markedly increased in the ovaries of *Esr2-/-* rats (Figure 1E-G). We further investigated the status of follicle activation in *Esr2* mutant rats prior to gonadotropin treatment.

### Loss of ESR2 leads to the activation of primordial follicles

We detected an increase in the recruitment of primordial follicles to the growing pool in PND28 *Esr2-/-* rat ovaries in the absence of any gonadotropin treatment (Figure 2A-C, J, M). To determine the postnatal period during which increased follicle activation becomes evident, further ovarian follicle counting was performed on PND16 and PND8. A decrease in primordial follicle counts with a corresponding increase in the number of activated follicles was detected in both PND16 (Figure 2D-F, K, N) and PND8 (Figure 2G-I, L, O) *Esr2-/-* ovaries. At all-time points, increased primordial follicle activation was found to be approximately twofold greater in *Esr2-/-* rats compared to wildtype. However, there was an increased number of atretic follicles in *Esr2-/-* ovaries (Figure 2B, E, H).

**Figure 2.**
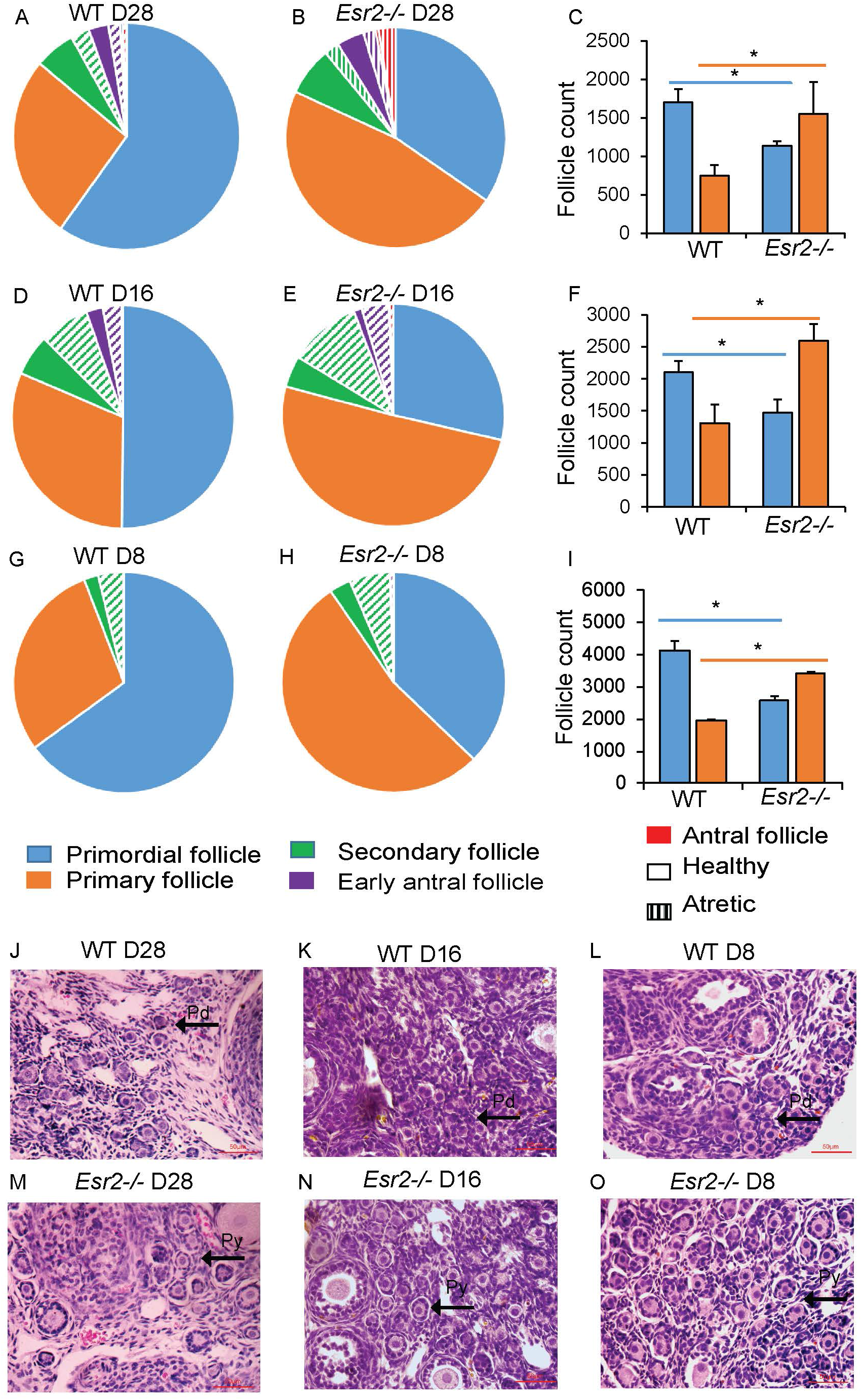
Increased activation of primordial follicles in *Esr2-/-* rat ovaries. Follicle counting in PND28 (**A-C, J, M**), PND16 (**D-F, K, N**), and PND8 (**G-I, L, O**) wildtype (WT) and *Esr2-/-* rats showed increased activation of primordial follicles in *Esr2-/-* ovaries. Primordial follicle activation was near two-fold starting on PND8 (**G-I, L, O**). A greater number of atretic follicles were present within the pool of activated follicles at PND28 (**A, B**) PND16 (**D, E**) and PND8 (**G, H**) in *Esr2-/-* ovaries. Data shown as mean ± SE, n≥3. \**P* ≤ 0.05. Pd, Primordial follicle; Py, Primary follicle.

### Loss of ESR2 results in premature ovarian senescence

We observed that total follicle counts in PND8 ovaries were similar between the wildtype and *Esr2-/-* rats; however, the primordial follicle counts were decreased by ~35% in *Esr2-/-* ovaries (Figure 3A). Further analyses of follicle count in 4wk, 12wk, and 24wk old *Esr2-/-* and age-matched wildtype rats revealed a sharp decline in primordial follicles in *Esr2-/-* ovaries (Figure 3B-H). Moreover, this decline in ovarian follicle numbers was associated with a significantly lower levels of serum estradiol and AMH in 24 wk old *Esr2-/-* rats. (Figure 3I, J).

**Figure 3.**
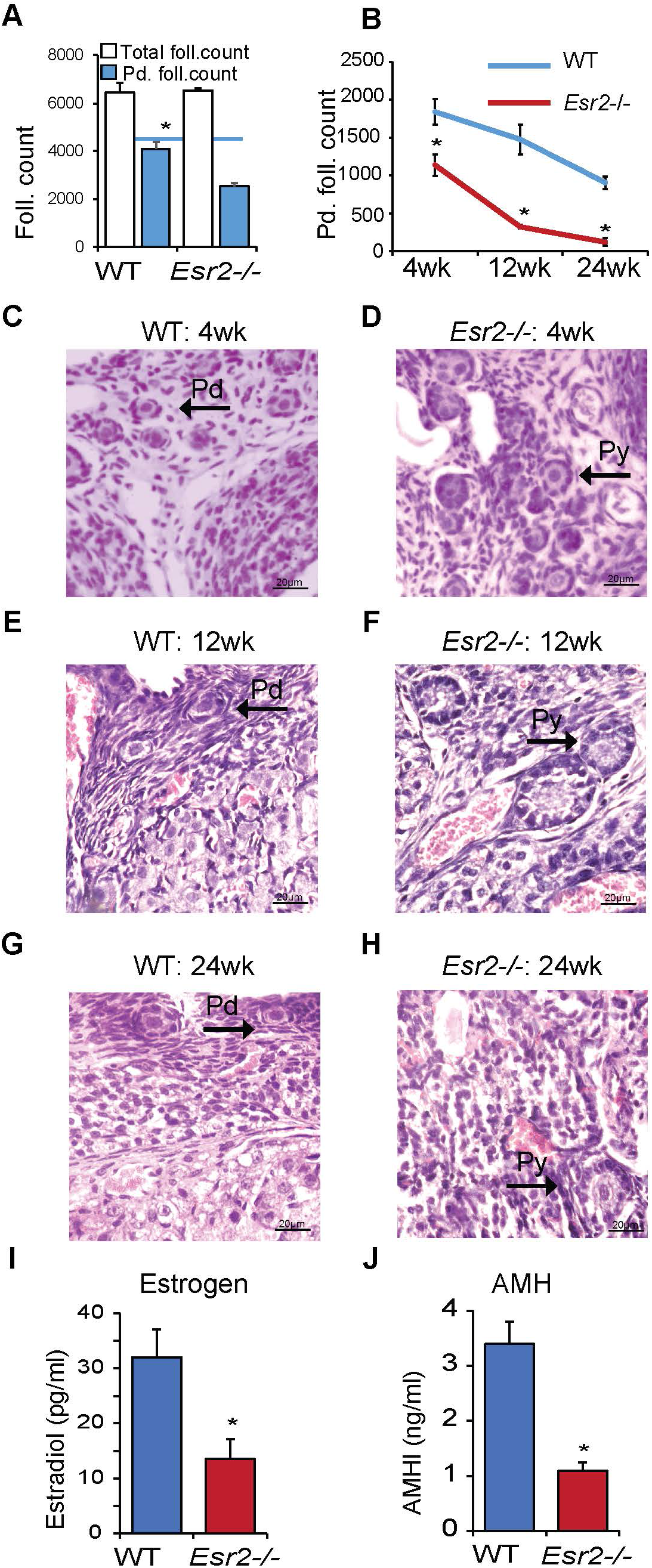
Premature ovarian senescence in *Esr2-/-* rats. The total number of follicles in PND8 ovaries were similar between wildtype (WT) and *Esr2-/-* rats; however, the number of primordial follicles were decreased in *Esr2-/-* ovaries (**A**). Follicle counts in 4wk, 12wk, and 24wk old WT and *Esr2-/-* rats revealed a sharp decline in the primordial follicle reserve in *Esr2-/-* ovaries (**B-H**). 24wk old *Esr2-/-* rats showed a significantly lower level of serum estradiol and AMH compared to WT (**I, J**). Data shown as mean± SE, n≥3 (follicle counting) and n≥6 (hormone assays), * *P*≤ 0.05. Pd, Primordial follicle; Py, Primary follicle.

### Regulation of primordial follicle activation is ESR2-dependent

Similar to *Esr2-/-* rats, an increased activation of primordial follicles was also observed in rats carrying a homozygous mutation in the DBD of *Esr2* (Figure 4A, B, G, I, J). The number of growing follicles was markedly increased while the number of primordial follicles decreased. However, this altered recruitment of primordial follicles was not detected in *Esr1*-/- rat ovaries (Figure 4A, C, G, I, K). In addition, treatment of wildtype rats with a selective ESR2-antagonist (PHTPP) resulted in increased activation of primordial follicles (Figure 4D, E, H, L, M). In contrast, treatment of wildtype rats with a selective ESR2-agonist (DPN) increased the proportion of follicles in the primordial state (Figure 4D, F, H, L, N).

**Figure 4.**
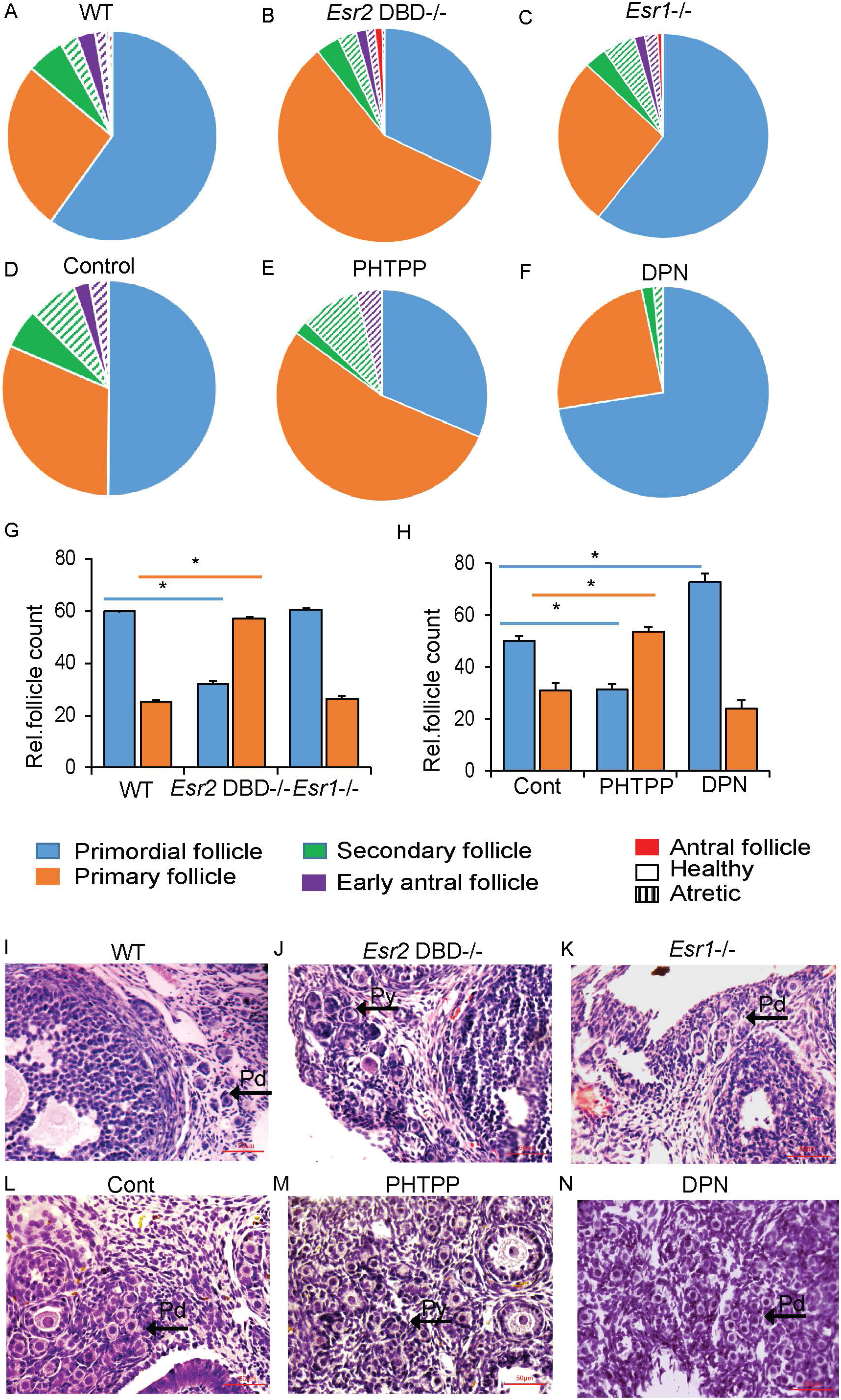
Regulation of primordial follicle activation is ESR2-dependent. Deletion of the ESR2 DBD increased primordial follicle activation at PND28 ovaries (**A, B, G, I, J**), but this was not observed in ovaries from Esr1-/- rats (**C, G, K**). Administration of a selective ESR2 antagonist, PHTPP, into wildtype rats increased primordial follicle activation (**D, E, H, L, M**), whereas treatment with an ESR2 agonist, DPN, suppressed the activation (**F, H, N**). Data shown as mean ± SE, n ≥ 3. * *P* ≤ 0.05. Rel., Relative. Pd, Primordial follicle; Py, Primary follicle.

### Disruption of ESR2 resulted in activation of the AKT and mTOR pathways

We detected abundant expression of ESR2 mRNA and protein in PND4, 6 and 8 ovaries (Figure 5 A, B). Expression of *Esr2* was also evident in primordial and primary follicles isolated from wildtype rat ovaries (Figure 5 C-E). Previous studies have shown that primordial follicle activation is regulated by the PI3K-PTEN-AKT and the mTORC1-TSC1/2-P70S6K signaling pathways (14,53). We therefore assessed the activation status of selected signaling molecules within these pathways in PND8 *Esr2-/-* ovaries and compared with that of wildtype. We observed increased activation of pAKT (T308 and S473) (Figure 5F-I), pERK1/2 (T202/Y204) (Figure 5L-M), mTORC1 (S2448) (Figure 5P-Q), and the downstream targets p70S6K (T389) (Figure 5R-S) and pRPS6 (S240/244) (Figure 5T-U). However, no significant alteration in PTEN levels (Figure 5J-K) or pTSC2 (T1462) phosphorylation (Figure 5N-O) was observed in *Esr2-/-* ovaries.

**Figure 5.**
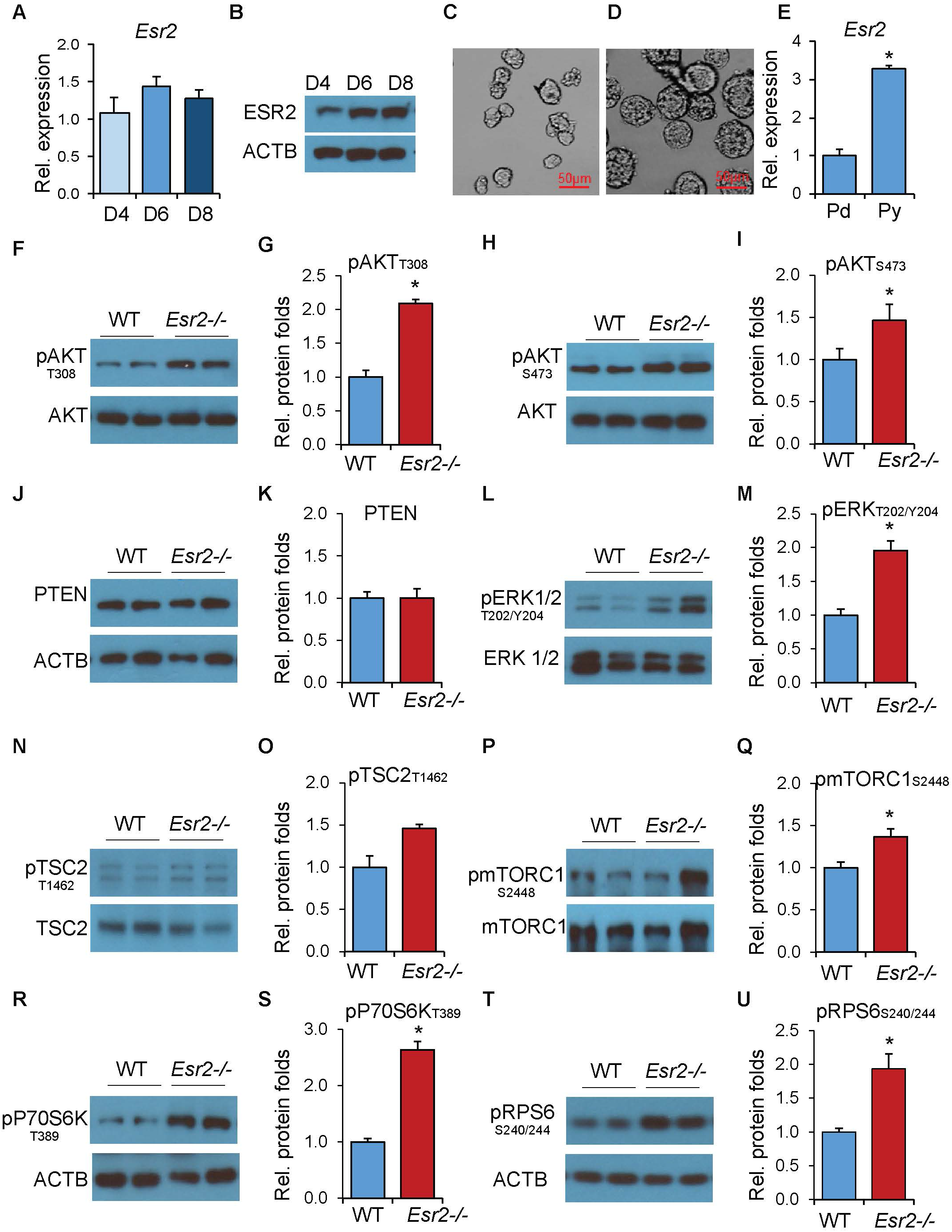
Activation of AKT, ERK, and mTOR signaling in *Esr2-/-* ovaries. Expression of ESR2 was detected in PND4, 6, and 8 rat ovaries using RT-qPCR (**A**) and western blotting (**B**). Primordial (Pd) (**C**), and primary (Py) (**D**) follicles were isolated from rat ovaries by digestion with liberase followed by size fractionation with strainers. RT-qPCR analysis demonstrated *Esr2* expression in both Pd and Py follicles (**E**). Western blot analyses of PND8 ovaries and quantification of signal intensities demonstrated a significant increase in AKT (**F-I**) and ERK1/2 (**L-M**) activation. This was associated with increased activation of mTORC1 (**P-Q**) and its targets P70S6K (**R-S**) and RPS6 (**T-U**). But no difference was observed in PTEN (**J-K**) and pTSC2 (**N-O**) levels. Signal quantification data are presented as mean ± SEM. n ≥ 6. \**P* ≤ 0.05. Rel., Relative.

### Loss of ESR2 increased expression of factors upstream of AKT and mTOR signaling

ESR2 is a transcriptional regulator and we have determined that the DB-dependent transcriptional activation function of ESR2 is required for regulation of primordial follicle activation (Figure 4). Therefore, we assessed the expression of factors that are known to activate the AKT and mTOR pathways and increase primordial follicle activation. Upstream regulators of AKT signaling including *Kitlg*, *Kit*, and *Igf1* (Figure 6A-C) as well as *Adcyap1r1* (Figure 6F) were upregulated in PND8 *Esr2-/-* ovaries. Oocyte derived TGFβ family members including *Bmp4, Bmp15,* and *Gdf9* were also significantly upregulated in *Esr2-/-* ovaries (Figure 6G-I). In addition, transcriptional regulators *Gata4* (Figure 6N), and *Npm2* (Figure 6O) were markedly upregulated in *Esr2-/-* ovaries. However, expression of *Fgf2* (Figure 6D), *Ntrk2* (Figure 6E), *Amh* (Figure 6J), *Bdnf* (Figure 6K), *Nobox* (Figure 6L), and *Sohlh1*(Figure 6M) was similar between *Esr2-/-* and wildtype ovaries.

**Figure 6.**
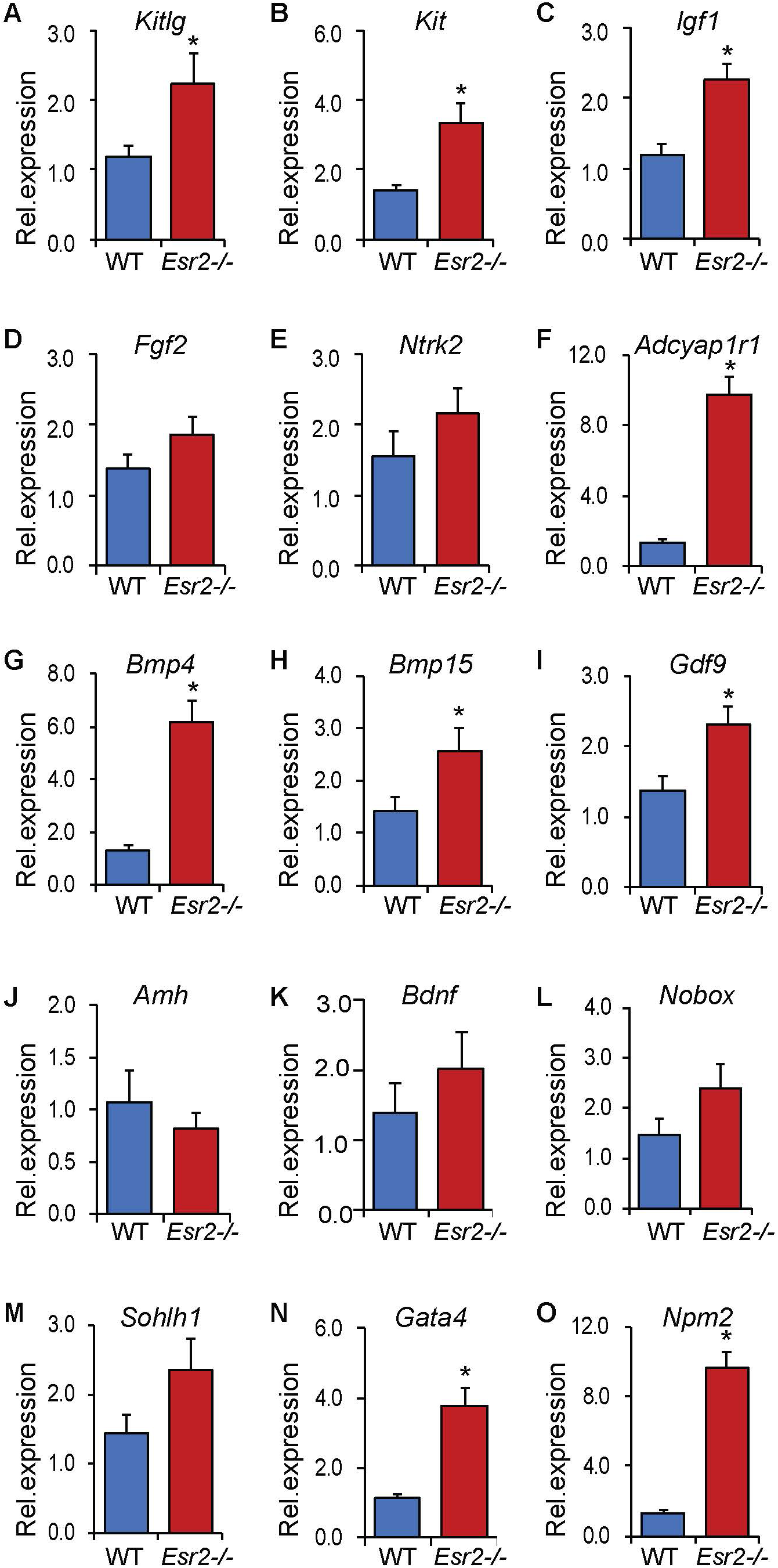
Transcript levels of known activators of the AKT and mTOR pathways. RT-qPCR was performed on PND8 wildtype (WT) and *Esr2-/-* rat ovaries to analyze the expression of genes involved in activation of the AKT and mTOR pathways and activation of primordial follicles. While the expression of *Kitlg* (**A**), *Kit* (**B**), *Igf1* (**C**), *Bmp15* (**H**) and *Gdf9* (**I**) was moderately upregulated in *Esr2-/-* rat ovaries, *Adcyap1r1*(**F**), *Bmp4* (**G**), *Gata4* (**N**), and *Npm2* (**O**) were highly upregulated. But no significant differences in the expression of *Fgf2* (**D**), *Ntrk2* (**E**), *Amh* (**J**), *Bdnf* (**K**), *Nobox* (**L**) and *Sohlh1*(**M**) were observed between the WT and *Esr2-/-* rat ovaries. RT-qPCR data represent the mean ± SEM. n ≥ 8. \**P* ≤ 0.05. Rel., Relative.

## Discussion

During the isolation of oocytes from the gonadotropin stimulated PND28 ovaries, we were surprised by a threefold higher yield in *Esr2-/-* rats. This appears to be due to the presence of a large number of activated follicles in *Esr2-/-* ovaries that were able to respond to gonadotropin treatment. This increased number of activated follicles was associated with a decreased number of primordial follicles suggesting excessive recruitment of primordial follicles. It is well known that activation of primordial follicles is independent of gonadotropin signaling (11). Thus, increased activation of primordial follicles observed in *Esr2-/-* rats was presumably independent of gonadotropin stimulation. It was confirmed by an increased activation of primordial follicles in PND8 and PND16 *Esr2-/-* ovaries and these findings suggest involvement of an intraovarian mechanism, which was defective during the early ovarian development due to the lack of ESR2.

Estrogen signaling plays a crucial role in ovarian follicle assembly, follicle development, and ovulation (42,43). Previous studies have shown that increased activation of ESR2-signaling can result in aberrant assembly of primordial follicles (42,43,54,55). However, we did not detect any significant changes in total follicle counts in PND8 ovaries due to the disruption of ESR2-signaling. Rather, we did observe a reduced number of primordial follicles and increased numbers of primary and other growing follicles. Our findings suggest that ESR2 functions in controlling the entry of primordial follicles into the growing pool, and thus plays a decisive role in determining female reproductive longevity.

In spite of normal follicle assembly and the establishment of the primordial follicle pool, excessive activation resulted in premature depletion of the primordial follicles in *Esr2-/-* rats. By 24 weeks of age, *Esr2-/-* rats contained less than 15% of the primordial follicles observed in wildtype ovaries (Fig 3B). As expected, the rapid decline in the follicle reserve led to premature ovarian senescence associated with a low AMH and estradiol level. AMH is an established indicator of follicle reserve and the serum level correlates well with the fertility potential (56,57). As polymorphisms or mutations can affect ESR2 function (35–38) and some endocrine disruptors can act as ESR2 agonists or antagonists (58–61), ESR2 regulation of primordial follicle activation can have a significant impact on women’s health and fertility.

Disruption of either ESR1 or ESR2 signaling results in dysregulation of gonadotropin secretion associated with a failure in follicle development and ovulation (39–41). Administration of exogenous gonadotropins fails to induce normal follicle maturation and ovulation in these mutant animals suggesting that disruption of estrogen signaling results in a primary ovarian defect (39–41). Increased activation of primordial follicles in *Esr2-/-* rats represents such a primary intraovarian defect. Absence of any increased activation of primordial follicles in *Esr1-/-* rats indicates that the regulatory mechanism is specific to ESR2. Furthermore, complete loss of ESR2 or loss of the DBD of ESR2 led to a similar phenotype, highlighting the requirement of the canonical transcriptional function of ESR2 in maintaining the primordial follicle reserve. In line with these observations, manipulation of ESR2 signaling by administration of selective antagonist or agonist into wildtype rats demonstrated that ESR2 plays a gatekeeping role during primordial follicle recruitment. Thus, transcriptional activation of ESR2 with selective ligands may prospectively increase the reproductive longevity in females. Selective ESR2-antagonists may also prove effective for *in vitro* activation (IVA) of primordial follicles.

It has been previously demonstrated that increased activation of AKT leads to accelerated activation of primordial follicles and depletion of the follicle reserve (62,63). Thus, increased activation of AKT in *Esr2-/-* rats may be related to the increased activation of primordial follicles. *Esr2-/-* rat ovaries also demonstrated increased activation of the mTOR and ERK pathways. Activated AKT can activate the mTOR pathway, which can also promote primordial follicle activation (14,53). A recent study has shown that ERK activation can induce primordial follicle activation through the KIT-mTOR signaling (64). Due to the complex architecture of the ovary (containing follicles at different stages and extrafollicular cells), it is difficult to discern whether these changes in signaling pathways are strictly due to accelerated primordial follicle development or reflect changes within primordial follicles.

While screening for candidate genes that may act as upstream regulators of the PI3K-AKT pathway, we identified upregulated expression of several growth factors and cytokines in *Esr2-/-* ovaries (Figure 6,7). Among these growth factors, KITLG and IGF1 can activate receptor tyrosine kinases on oocytes and induce activation of primordial follicles (22,65–67). The other upregulated factors are oocyte-derived TGFβ family members including BMP4, BMP15, and GDF9, which are known to stimulate follicle activation (19,20). While GDF9 and BMP15 can activate the mTOR pathway (68), BMP signaling can also activate the ERK pathway through TAK1 (69). In addition to these factors, upregulation of ADCYAP1R1, and NPM2 were prominent in PND8 *Esr2-/-* rat ovaries, which might have also contributed to the activation of the PI3K-AKT pathway. NPM2 is a transcriptional regulator in oocytes (70), which was found to activate the AKT (71,72), mTOR, and ERK pathways in other cells (73). While ADCY3 is expressed in oocytes, its receptor ADCYAP1R1 is expressed in GCs; ADCY3 signaling generates cAMP, which in turn can activate the AKT pathway (74). As stated previously, these factors may regulate follicle activation but how ESR2 is implicating them in the ovary is currently unclear.

**Figure 7.**
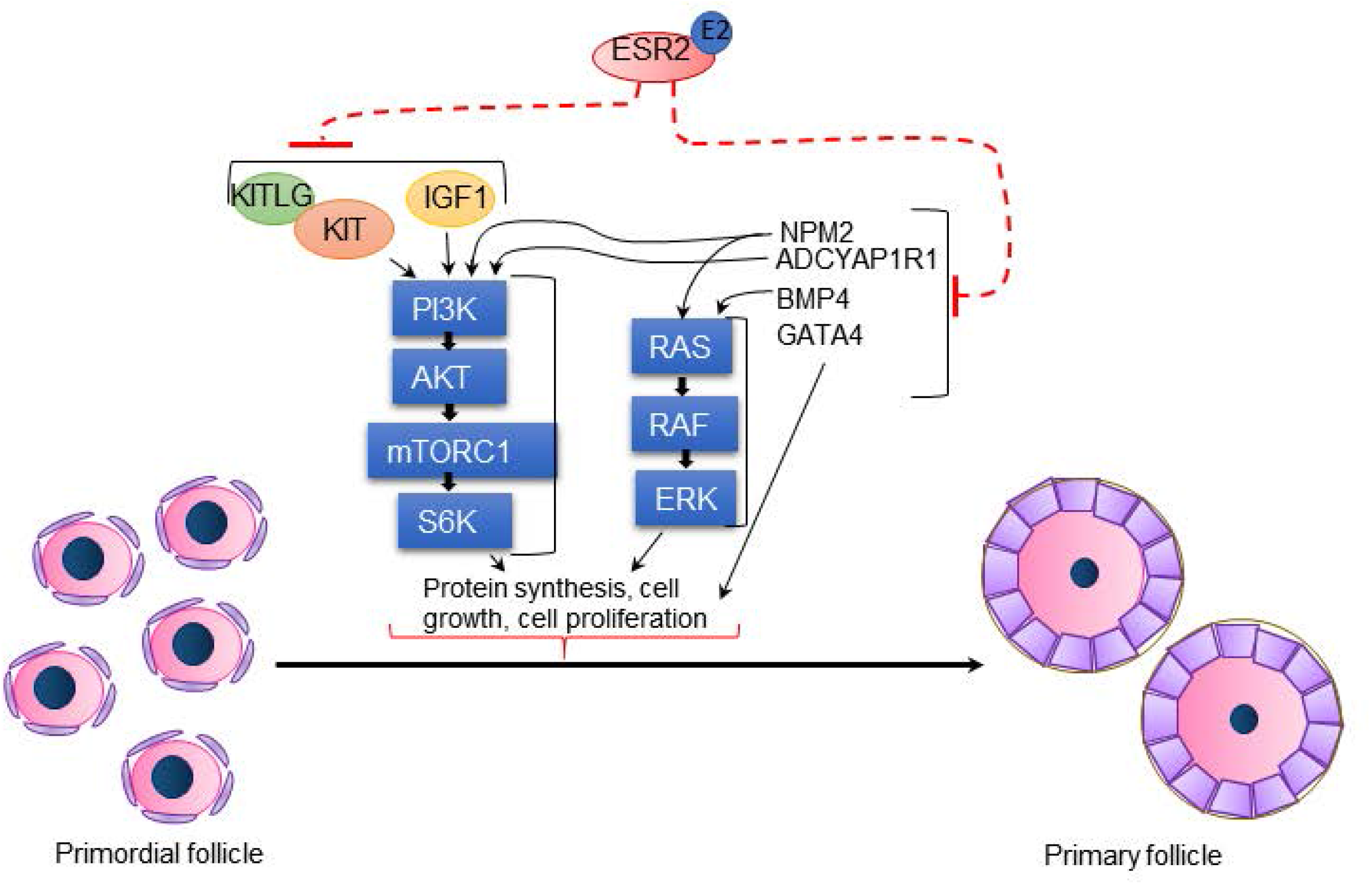
ESR2 signaling in the regulation of primordial follicle activation. Loss of ESR2 leads to upregulation of both granulosa cell and oocyte derived factors that can activate the AKT, ERK and mTOR pathways. Increased levels of KITLG, KIT and IGF1, as well as NPM2, and ADCYAP1R1 can activate the AKT pathway followed by the mTOR pathway. Moreover, upregulation of NPM2 and BMP4 can activate the ERK pathway. Activated AKT, ERK and mTOR pathways in association with the transcriptional regulator GATA4 can promote the transition of primordial follicles to primary follicles in *Esr2-/-* ovaries.

In this study, we demonstrate a novel role of ESR2 in maintaining the primordial follicle reserve. Excessive activation of primordial follicles leads to premature loss of the follicle reserve. Our findings suggest that ESR2 is involved in downregulating the expression of ovarian-derived factors, which function to activate key signaling pathways that induce primordial follicle activation. Loss of ESR2 upregulates these factors and results in an increased activation of the signaling pathways that promotes primordial follicle activation.

## Acknowledgments

We acknowledge the financial support from the KUMC School of Medicine (SOM Bridging Grants) for completion of this research.

## Grants or fellowships support

This study was supported in part by the School of Medicine Bridging grant from the University of Kansas Medical Center.

